# Comparative Analysis of Machine Learning Algorithms for Breast Cancer Classification: SVM Outperforms XGBoost, CNN, RNN, and Others

**DOI:** 10.1101/2024.04.22.590658

**Authors:** Prithwish Ghosh, Debashis Chatterjee

## Abstract

This study evaluates ten machine learning algorithms for classifying breast cancer cases as malignant or benign based on physical attributes. Algorithms tested include XGBoost, CNN, RNN, AdaBoost, Adaptive Decision Learner, fLSTM, GRU, Random Forest, SVM, and Logistic Regression. Using a robust dataset from UCI machine learning Breast Cancer, SVM emerged as the most accurate, achieving 98.2456% accuracy. While AdaBoost, Logistic Regression, Neural Networks, and Random Forest showed promise, none matched SVM’s accuracy. These findings underscore the potential of machine learning, particularly SVMs, in cancer diagnosis and treatment by analyzing physical attributes for improved diagnostics and targeted therapies.

## INTRODUCTION

Breast cancer is a type of cancer that develops in the cells of the breast. It is one of the most common cancers among women worldwide, but it can also affect men, though it’s rare. Early detection through screening, such as mammograms, and advances in treatment have significantly improved the prognosis for many people diagnosed with breast cancer [9]. Treatment options typically include surgery, chemotherapy, radiation therapy, hormone therapy, targeted therapy, or a combination of these approaches, depending on the type and stage of the cancer [30].

Machine learning (ML) algorithms have revolutionized various scientific fields in recent years. [36] developed a Computer-Aided Diagnosis (CAD) system using Machine Learning (ML) and region-growing segmentation to analyze breast ultrasound images. [2] proposed using B-mode and elastography images for breast cancer detection. Their system utilized 82 ultrasound images, employing geometrical and texture features. [8] developed a CAD system based on morphological features from B-mode ultrasound images. [11] proposed a CAD system employing various classifiers to classify breast ultrasound images based on textures and morphological features, with Linear Discriminant Analysis (LDA) performing best [20].

Other researchers like [33], [17], [25], and [19] introduced different approaches using SVM, LDA, and Modified Neural Network (MNN) achieving notable accuracies ranging from 75.94% to 97.80%. Additionally, methods by [14] using logistic regression, [4] utilizing XGBoost, and [10] employing morphological features showed promising results with accuracies around 89.40% to 94.0%. Lastly, [15] proposed a deep learning approach combining semantic segmentation and DenseNet201 with SVM.

### Objective of the Paper

This study takes aim at a critical question: which machine learning approach reigns supreme in classifying breast cancer based on physical attributes? We intend to assess the effectiveness of machine learning, Neural Networking, and Deep Learning techniques in predicting Breast Cancer Classification based on class. We unleashed a diverse arsenal of ten classification methods, including neural networks and deep learning algorithms, on a robust dataset. This finding underscores the crucial role of choosing the right tool for the job in breast cancer diagnosis. By strategically implementing these techniques, we achieved near-perfect accuracy, a significant leap forward compared to traditional machine learning methods.

## THE DATASET

The dataset of breast cancer [34]. Characteristics are derived from a digitized image of a breast mass’s fine needle aspirate (FNA), depicting attributes of the cell nuclei within the image. Some sample images can be accessed at http://www.cs.wisc.edu/~street/images/.

### Attribute Information

ID number, Diagnosis (M = malignant, B = benign), Ten real-valued features are computed for each cell nucleus:, radius (mean of distances from the center to points on the perimeter), texture (standard deviation of gray-scale values), perimeter, area, smoothness (local variation in radius lengths), compactness (perimeter^2^ / area - 1.0), concavity (severity of concave portions of the contour), concave points (number of concave portions of the contour), symmetry, fractal dimension

## METHODOLOGIES

This study embarked on a mission to identify the most effective machine learning warrior in the fight against breast cancer. We assembled an arsenal of ten classification algorithms, each a powerful tool for analyzing physical attributes and distinguishing between malignant and benign tumors.

For all the algorithms we choose *X* = [*x*_1_, *x*_2_, …, *x*_*n*_] which represent the input features, where each *x*_*i*_ represents a vector of features including the parameters: Radius (mean of distances from the center to points on the perimeter): 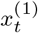, Texture (standard deviation of gray-scale values): 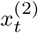, Perimeter: 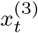, Area: 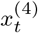, Smoothness (local variation in radius lengths): 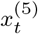, Compactness (perimeter^2^ / area - 1.0): 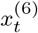, Concavity (severity of concave portions of the contour): 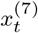, Concave points (number of concave portions of the contour): 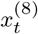, Symmetry: 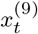, Fractal dimension (“coastline approximation” - 1): 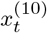

Let *y* denote the target variable, which is the Diagnosis (Malignant or Benign)

### The Algorithm Legion

Our ten valiant contenders included:

I. Support Vector Machine (SVM) [31]: A veteran classifier known for its ability to find clear boundaries between data-points [13].
II. Random Forest: A committee-based approach that leverages the wisdom of multiple decision trees for robust predictions [3] [23].
III. Logistic Regression: A workhorse algorithm that calculates the probability of an outcome based on its features [35] [29].
IV. XGBoost is a popular gradient boosting algorithm known for its efficiency and performance in solving regression, classification, and ranking problems by sequentially building a series of decision trees. [22, 9] [5, 21]
V. AdaBoost: A champion for boosting the performance of weaker learners by strategically focusing on challenging data points [27] [12, 7].
VI. Adaptive Decision Learner: A dynamic approach that tailors decision trees to the specific characteristics of the data.
VII. Neural Network Variants: We utilized the power of three neural network architectures [1] [32]:
  a. Convolutional Neural Network (CNN): An expert at identifying patterns in image data, even if they’re subtly hidden [28].
  b. Long Short-Term Memory (LSTM): A master at handling sequential data, potentially useful for capturing the progression of the disease [18].
  c. Gated Recurrent Unit (GRU): Another sequential data specialist, offering an alternative approach to LSTMs [6] [24].
  d. Recurrent Neural Networks (RNNs) are a class of artificial neural networks designed to efficiently process sequential data by maintaining an internal state, allowing them to capture temporal dependencies within the input sequences. [26] [18][16]

## RESULTS

Our research utilized ten distinct machine learning algorithms to discern Breast Cancer cases (Malignant or Benign), relying on their physical attributes. The algorithms employed in our investigation mentioned in the section

We utilized real-valued features for each cell nucleus: radius (mean of distances from the center to points on the perimeter), texture (standard deviation of gray-scale values), perimeter, area, smoothness (local variation in radius lengths), compactness (perimeter^2^ / area - 1.0), concavity (severity of concave portions of the contour), concave points (number of concave portions of the contour), and symmetry. The target variable was Diagnosis (Malignant, Benign).

After rigorous training and testing, the Support Vector Machine Classifier exhibited the highest accuracy among all the algorithms, with the Random Forest Classifier closely following with an accuracy of 98.24%, surpassing the other classifiers in predictive performance.

This outcome indicates that the Random Forest Classifier is notably effective for Breast Cancer classification based on physical characteristics. The ensemble nature of the SVC algorithm likely contributed to its superior performance in capturing intricate relationships within the data.

This has significant implications for various medical or biological studies and can aid in understanding the properties and characteristics of cancer cells in bodies.

The accuracy scores for each algorithm are presented in Table 1, where it is evident that the SVC Classifier achieved the highest accuracy at 98.2456%. Other algorithms, such as AdaBoost, Logistic Regression, Neural Networks, and Random Forest, also demonstrated commendable accuracy rates but were outperformed by the SVC Classifier. The remaining algorithms, including CNN, GRU, RNN, XGBoost, and LSTM, exhibited reasonable performance, albeit with slightly lower accuracy scores than the top-performing algorithms, as depicted in Figure 2.

**Table 1.** By using the ten different Deep Learning and machine learning algorithms on the data after missing value imputations, we get that the Random Forest Classifier Algorithm gives us the best result among them per the accuracy score. All of the accuracy scores are mentioned in the table.

| ML Algorithm | Accuracy of the algorithm |
| --- | --- |
| XGboost | 95.6140 |
| Neural Networks | 97.368 |
| CNN | 94.736 |
| RNN | 94.7 |
| AdaBoost | 97.368 |
| Adaptive Decision Learner | 93.859 |
| LSTM networks | 94.736 |
| Gated Recurrent Units | 93.859 |
| Random Forest Classifier | 96.4912 |
| Support Vector Machine | 98.2456 |
| Logistic Regression | 97.368 |

**Fig. 1.**
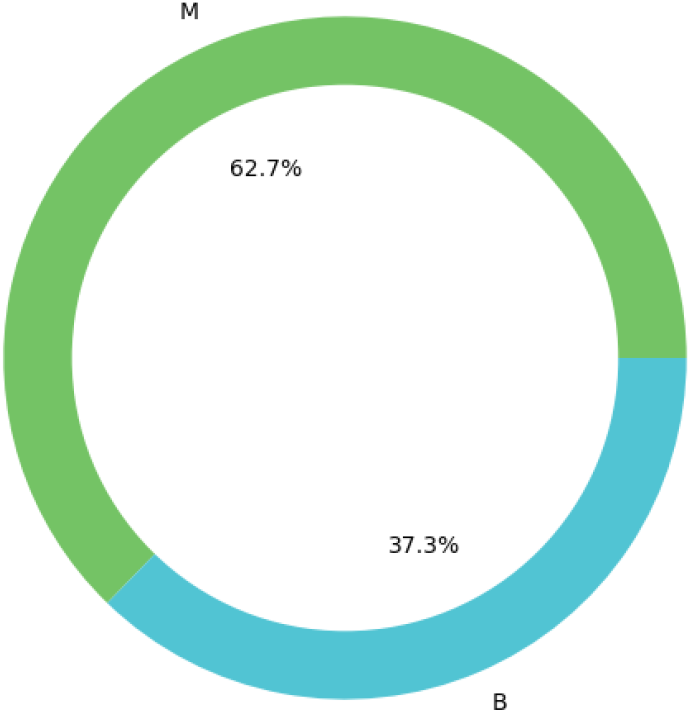
A cylindrical plot concerning Breast Cancer Classification where we can say that we have 62.7% Maligant and 37.3% Benign from our data.

**Fig. 2.**
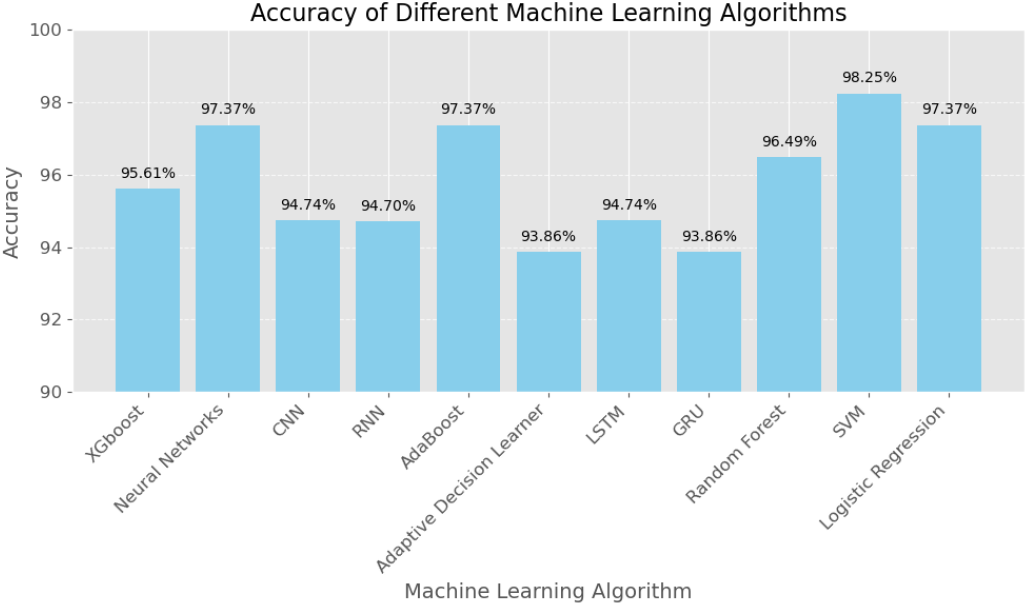
Using the 11 different machine learning algorithms, we get the best result from the SVM Classifier Algorithm. A pictorial bar diagram concerning their accuracy score for 11 different Deep and Machine learning methods is given in this plot.

## CONCLUSION

The results section concludes that the Support Vector Machine (SVM) Classifier achieved the highest accuracy at 98.2456%, making it the most effective algorithm for categorizing Breast Cancer cases based on physical attributes. While other algorithms like AdaBoost, Logistic Regression, Neural Networks, and Random Forest also showed reasonable accuracy rates, they were surpassed by the SVM Classifier. The remaining algorithms, including CNN, GRU, RNN, XGBoost, and LSTM, demonstrated reasonable performance but slightly lower accuracy than the top-performing algorithms. Overall, these findings underscore the efficacy of machine learning algorithms, particularly the SVM Classifier, in accurately categorizing Breast Cancer data, which has significant implications for medical and biological studies, aiding in understanding the properties and characteristics of cancer cells within the body.

## Author contributions statement

D.C. designed, conceptualized, and developed the research and synthesized interdisciplinary statistical methodologies and models. P.G. conceptualized the model, collected and prepared the datasets, wrote codes for various modified datasets, and performed code-based analysis (mainly using Python). P.G., D.C., wrote, modified, and reviewed the manuscript.

## Data Availability

The Breast Cancer Classification data of [34] that we have used in this study is publicly available from the GitHub Machine Learning Repository [http://www.cs.wisc.edu/~street/images/]

## Code Availability Statement

The code generated in this paper’s results is publicly available on GitHub at [https://github.com/Prithwish-ghosh/Breast-Cancer].

We have provided the code under the public license, allowing researchers to reproduce our results and facilitate further development in this area.

